# Developmental Effects on Relative Use of PEPCK and NADP-ME Pathways of C_4_ Photosynthesis in Maize

**DOI:** 10.1101/2021.06.25.449949

**Authors:** Jennifer J. Arp, Shrikaar Kambhampati, Kevin L. Chu, Somnath Koley, Lauren M. Jenkins, Todd C. Mockler, Doug K. Allen

## Abstract

C_4_ photosynthesis is an adaptive photosynthetic pathway which concentrates CO_2_ around Rubisco in specialized bundle sheath cells to reduce photorespiration. Historically, the pathway has been characterized into three different subtypes based on the decarboxylase involved, although recent work has provided evidence that some plants can use multiple decarboxylases, with maize in particular using both the NADP-malic enzyme (NADP-ME) pathway and phosphoenolpyruvate carboxykinase (PEPCK) pathway. Parallel C_4_ pathways could be advantageous in balancing energy and reducing equivalents between bundle sheath and mesophyll cells, in decreasing the size of the metabolite gradients between cells and may better accommodate changing environmental conditions or source to sink demands on growth. The enzyme activity of C_4_ decarboxylases can fluctuate with different stages of leaf development, but it remains unclear if the pathway flexibility is an innate aspect of leaf development or an adaptation to the leaf microenvironment that is regulated by the plant. In this study, variation in the two C_4_ pathways in maize were characterized at nine plant ages throughout the life cycle. Two positions in the canopy were examined for variation in physiology, gene expression, metabolite concentration, and enzyme activity, with particular interest in asparagine as a potential regulator of C_4_ decarboxylase activity. Variation in C_4_ and C_3_ metabolism was observed for both leaf age and canopy position, reflecting the ability of C_4_ pathways to adapt to changing microenvironments.

**One Sentence Summary:** The proportion of the two C_4_ pathways in maize plants is dependent on canopy position and not the age of the leaf.

## Introduction

C_4_ photosynthesis is a beneficial adaptation to environmental conditions that enables plants to more effectively assimilate carbon dioxide relative to C_3_ plants, resulting in some of the most productive crops on a biomass basis. C_4_ photosynthesis has evolved over 60 times in plants to overcome the inefficiencies of Rubisco by minimizing the oxidation reaction and photorespiration (Sage et al., 2011). The pathway has historically been categorized into three subtypes based on the decarboxylase—NAD-malic enzyme (NAD-ME), NADP-malic enzyme (NADP-ME) or phospho*enol*pyruvate carboxykinase (PEPCK)—but additional evidence has identified plants that can use a combination of decarboxylases in C_4_ photosynthesis (Chapman and Hatch, 1981; Furbank, 2011; Wang et al., 2014). Several advantages of using a combination of C_4_ pathways have been proposed. By dividing the transfer of metabolites between the mesophyll and bundle sheath into a combination of amino and organic acids, smaller gradients of each are sufficient to drive the transfer between cell types (Pick et al., 2011; Stitt and Zhu, 2014; Wang et al., 2014). The two pathways distribute energy and reducing equivalents differently between the two cell types which could be complementary and provide faster response to shifting light environments (Furbank, 2011; Stitt and Zhu, 2014). The ratio of NADP-ME to PEPCK pathway flux may not be fixed within species, across plant development, or under changing environment. Literature reports in maize indicate PEPCK flux is 10-25% of total C_4_ flux (Chapman and Hatch, 1981; Weissmann et al., 2016; Arrivault et al., 2017), whereas in the C_4_ dicot *Flaveria bidentis,* 50% of assimilated carbon comes through the PEPCK pathway (Meister et al., 1996). More broadly, enzyme activities for key C_4_ enzymes in NADP-ME-type species also indicate extensive variation in the ratio of aspartate to malate translocated in C_4_ plants (Kanai and Edwards, 1999). The ratio of malate and aspartate can also fluctuate in response to nitrogen limitation (Khamis et al., 1992). Though multiple studies have observed differences in pathway flux, the factors that elicit particular C_4_ photosynthetic subtype use remain to be firmly established and could aid efforts to sustainably increase crop productivity.

The role for dual C_4_ pathways has been considered over plant development in *Cleome gyandra* (Sommer et al., 2012). In this dicot species, decarboxylase activity varied with leaf age along the canopy using the NAD-malic enzyme pathway and a supplemental PEPCK pathway. In younger leaves, the activity of NAD-ME was twice that of PEPCK; however, in older leaves, PEPCK activity increased and NAD-ME activity decreased, leading to nearly double PEPCK activity compared to NAD-ME while the total amount of decarboxylase activity remained nearly constant. Though the study was intended to characterize the establishment of C_4_ photosynthesis across development, the findings established within-plant differences and indicated the ratio of pathway use is pliable within species.

Canopy position and the resulting light environment of the leaf has an effect on the rate of photosynthesis. In the maize canopy, all new leaves form in full sunlight at the top of the canopy and become progressively shaded by new growth, requiring leaves to adapt to the new environment after they are fully formed. In the maize canopy, the gradients of light and age generate a pattern of increasing photosynthetic capacity at the top of the canopy, and less activity at the bottom of the canopy from older, self-shaded leaves (Ubierna et al., 2013; Niinemets, 2016; Pons, 2016; Collison et al., 2020). Up to 50% of maize photosynthesis occurs in these shaded parts of the canopy (Baker et al., 1988), and leaves in the lower canopy are capable of high rates of photosynthesis when not exposed to large degrees of self-shading (Collison et al., 2020). In addition to differences in overall rates of photosynthesis, foundational work on light regulation in maize has shown that the C_4_ enzymes are more light-controlled than the C_3_ enzymes (Sugiyama et al., 1984; Ward and Woolhouse, 1986). However, at low light, the C_3_ pathways are expected to be more downregulated than C_4_ as the plant shifts N away from Rubisco and toward light harvesting (Boardman, 1977; Björkman, 1981; Hikosaka and Terashima, 1995; Evans and Poorter, 2001; Walters, 2005; Tazoe et al., 2006; Pengelly et al., 2010). Moreover, species utilizing different C_4_ subtypes have different degrees of shade acclimation, with C_4_ grasses which use the NADP-ME subtype, such as *Z. mays*, able to acclimate to shade more readily than those using NAD-ME or PEP-CK subtypes (Sonawane et al., 2018).

The interaction between C_4_ pathways and plant nitrogen status is well-documented (Khamis et al., 1992; Taub and Lerdau, 2000; Ghannoum et al., 2005; Pinto et al., 2014; Pinto et al., 2016) but mechanistically is not completely understood. The PEPCK pathway shuttles carbon using the amino acid aspartate which provides a potential regulatory link between C_4_ photosynthesis and the nitrogen status of the plant. Aspartate can be rapidly converted to asparagine that is associated with important developmental cues during the maize life cycle (Seebauer et al., 2004) and transports carbon and nitrogen between leaves and developing kernels (Cañas et al., 2010). Asparagine is believed to play a role in signaling seed storage protein deposition (Hernández-Sebastià et al., 2005; Pandurangan et al., 2012) and, importantly, appears to be a regulator of PEPCK activity in seeds. When asparagine was added exogenously to grape seeds, PEPCK activity was increased 100-fold (Walker et al., 1997). This relationship is likely important *in vivo* because PEPCK proteins are developmentally regulated in tomato and grape seeds (Walker et al., 1999; Bahrami et al., 2001), and PEPCK activity coincides with peak amino acid metabolism and storage protein deposition, possibly through asparaginase activity (Walker et al., 1999). Despite known anaplerotic functional relationships between asparagine and PEPCK, it remains unclear if asparagine could be important for the regulation of the C_4_ PEPCK gene during development and aging of the maize leaf.

In this study, we grew maize plants at ten day intervals to sample leaves from the top and middle of the canopy at nine plant ages to determine the effect of canopy position and plant age on the proportion of C_4_ flux through PEPCK relative to NADP-ME. Parts of the canopy with the greatest fluctuations in light environment, including lower leaves, would be expected to use proportionally more mixed C_4_ pathways to allow leaf cells to maintain an energetic homeostasis. Additionally, plant age might affect the C_4_ pathways such that older plants would use proportionately more equal C_4_ pathways to maintain moderate levels of photosynthesis as leaf N is remobilized to the grain during senescence. One possible regulator of PEPCK variation based on work in other species is asparagine, so asparagine concentration across development was compared with C_4_ pathway activity to identify potential regulatory relationships. Overall, we found the plant modifies the proportion of the two C_4_ photosynthetic pathways in response to the developmental and environmental microenvironment of the leaf throughout the growing season.

## Results

### Distinguishing maize plant developmental progression from canopy influence

To understand variation in the C_4_ pathways over the course of the plant life cycle and within the canopy, maize plants (*var.* W22) were grown in ten-day intervals for 100 days in the greenhouse from February to July 2019. Plants were sampled on July 2-3, 2019, for the nine developmental times, representing plants 15 days after sowing to 100 days after sowing (DAS; Figure 1). Measurements were taken from the top collared leaf (“top leaf”) and leaf 13 (“subtending leaf”), which is the leaf subtending the ear in W22. All measurements were taken from the middle of the leaf, 20 cm from the leaf tip. The top leaf represents the leaf receiving the highest amount of sunlight and is the youngest expanded leaf on the maize plant. The subtending leaf provides the most nutrients to support ear growth (Subedi and Ma, 2005). Measurements were not taken from leaves within the whorl nor from leaves which had fully senesced. In total, these leaf samples represented the span of the maize life cycle and two functionally distinct parts of the leaf canopy.

**Figure 1:**
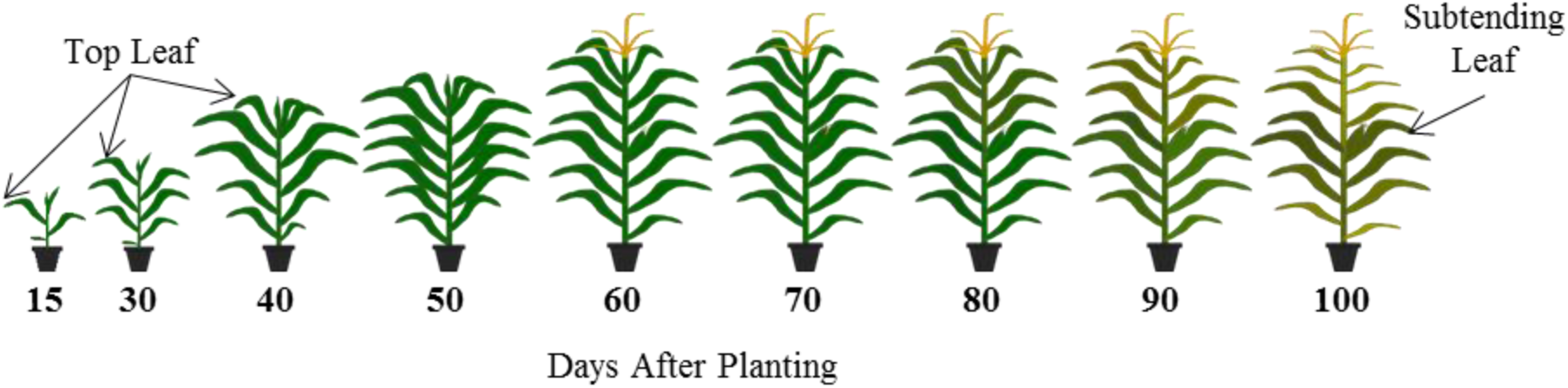
Experimental Design: Plants were grown at ten day intervals in the greenhouse to generate a gradient of maize development which could be sampled on the same day. Nine individual time points were used and captured maize development from the V3 stage until physiological maturity. The leaf at the top of the canopy for each time point was sampled until the top leaf senesced at 90 days after sowing. Also sampled was leaf 13, which subtends the ear in genotype W22. Leaf 13 emerged from the whorl at 50 days after sowing.

### Photosynthetic CO_2_ assimilation decreased with plant age

Overall photosynthetic rate has been shown to be variable during plant growth and in the canopy (Dwyer and Stewart, 1986; Chen et al., 2016; Niinemets, 2016), so we measured net CO_2_ assimilation (A_net_) on the plants at the nine growth stages. A_net_ was measured at both ambient (350 μmol m^−2^ s^−1^) and high (1500 μmol m^−2^ s^−1^) light levels in the LI-6800 chamber. During plant growth A_net_ consistently decreased with age in the plant for both top and subtending leaves at the ambient and high light levels (Figure 2A). Leaf position did not have a significant effect on A_net_ (p=0.90, ANOVA type II). The interaction between light and plant age was significant (p=5.96e-6, ANOVA): in young plants, A_net_ increased substantially in the high light conditions compared to 500 μmol m^−2^ s^−1^; however, the response to high light was dampened in older plants.

**Figure 2:**
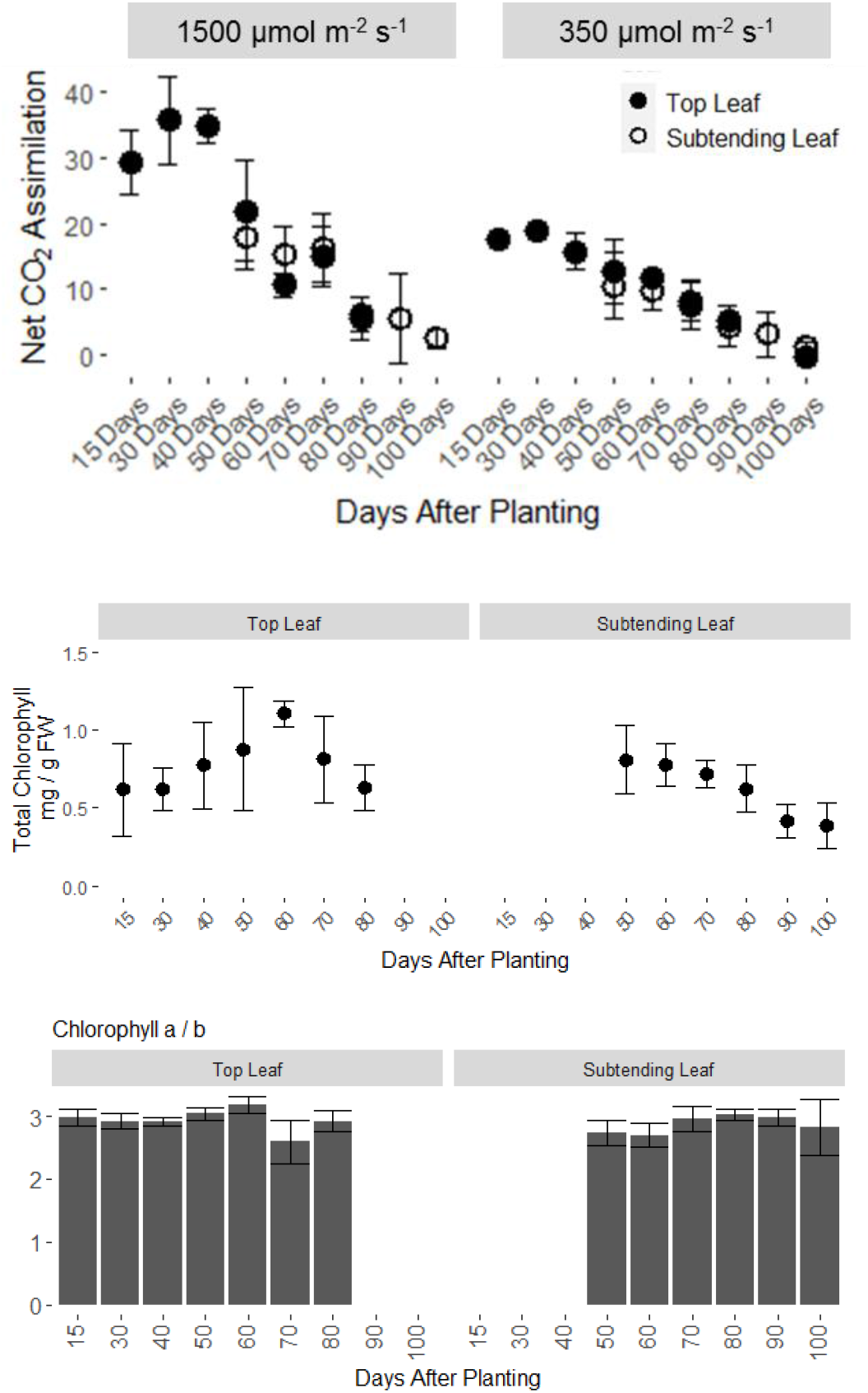
Measures of overall photosynthesis: A.) Net CO_2_ assimilation (A) was measured at two light levels using a LI-COR 6800 gas exchange instrument. B.) Total chlorophyll was measured using a spectrometric assay. C.) Chlorophyll a to b ratio. All data are means ± sd of 3-4 replicate plants.

### Chlorophyll content and A_net_ are correlated with plant age

A_net_ was modestly correlated with chlorophyll concentration in the leaf (Pearson’s R = 0.39) as an overall indicator of photosynthetic capacity. Total chlorophyll concentration increased in the top leaf from 0.6 mg chlorophyll g^−1^ FW to 1.1 mg g^−1^ FW between 15 DAS and 60 DAS before decreasing back to 0.6 mg g^−1^ FW at 80DAS, the last sampling time for the top leaf before complete senescence (Figure 2B). Leaf position in the canopy did not have a significant effect on total chlorophyll concentration (p=0.12, ANOVA type II); however, plant age had a significant effect (p=0.004). Chlorophyll concentration decreased steadily in the subtending leaf, likely as a consequence of the initiation of leaf senescence (Figure 2B). Throughout the growth cycle, the nitrogen status of the leaf and the light environment are expected to vary and can affect the chlorophyll a to b ratio (Hikosaka and Terashima, 1995). Chlorophyll a to b ratio was not correlated with the age of the plant or canopy position (p=0.14 and p=0.73, ANOVA type II); however, the interaction effect between leaf position and plant age was significant (p=0.001, ANOVA type II; Figure 2C), and the ratio drops slightly in the top leaves at 70 and 80 days after sowing, reflecting the beginning of senescence.

### Ratio of C_4_ decarboxylases changed with canopy position and plant age

The responses of the two C_4_ pathway decarboxylases were investigated using transcriptional analysis since the enzymes involved in the C_4_ pathway are at least partially regulated at this level (Pick et al., 2011). Using the gene expression atlas resource for maize (Stelpflug et al., 2016), gene expression levels were probed for the two decarboxylase enzymes in maize leaves over developmental age. Both PEPCK1 (GRMZM2G001696) and NADP-ME (NADP-ME3, GRMZM2G085019) expression varied with growth stage, beginning with the highest expression in the youngest leaves and decreasing with leaf age. NADP-ME3 expression decreased quickly after the V9 growth stage; its expression was approximately 2-fold higher than PEPCK1 until the V9 growth stage, where expression became approximately equal for the two decarboxylases (Figure 3A). Based on the transcriptomic evaluation, a comprehensive experiment was designed to assess the C_4_ subtypes through the lifetime of leaves.

**Figure 3:**
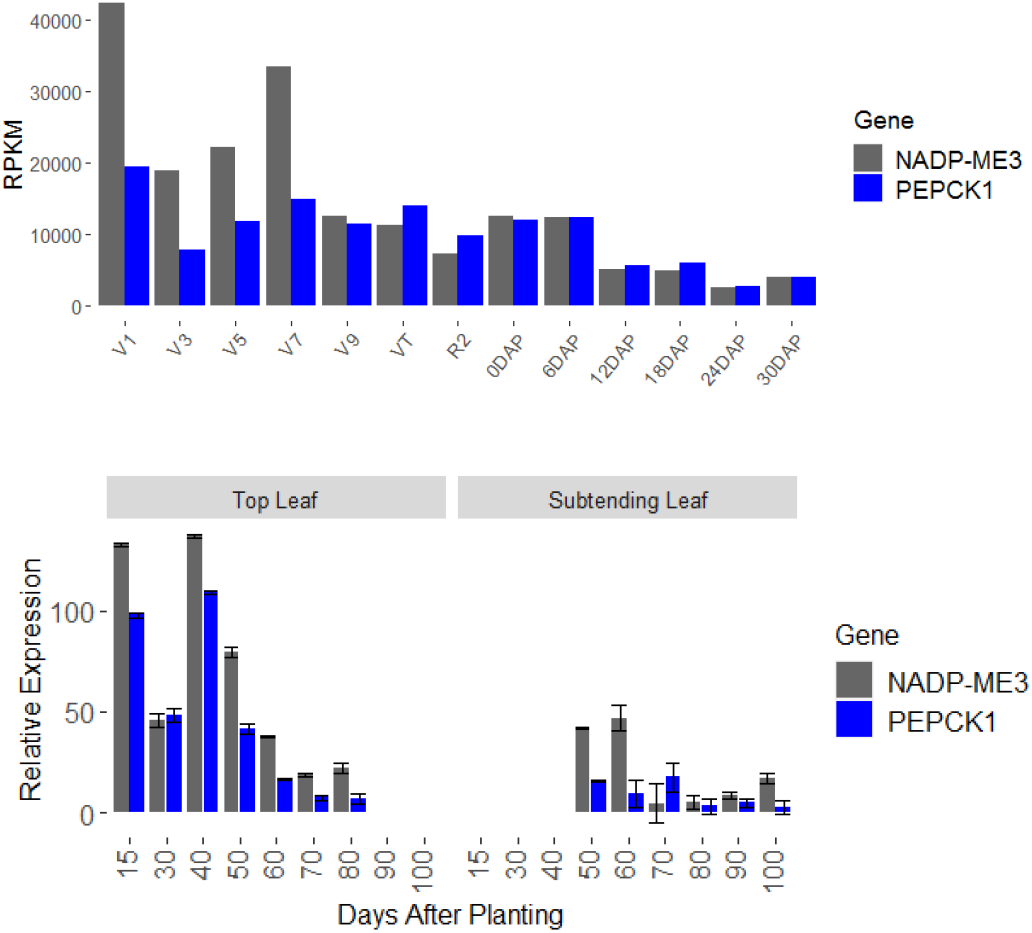
Gene expression differences: A.) Expression of NADP-ME3 (Zm00004b015828) and PEPCK1 (Zm00004b001002) in the maize gene expression atlas (Stelpflug et al, 2016) decreased with plant age, and the ratio of NADP-ME3 (gray) to PEPCK1 (blue) decreased in older plants. B.) Relative gene expression for PEPCK1 and NADP-ME3 was quantified using qPCR. Data are means ± sd of 3-4 replicate plants.

The expression of NADP-ME3 and PEPCK1 in the plants grown in the greenhouse was assessed by quantitative real-time PCR to confirm the trend indicated by the maize atlas. Similar to the pattern from the maize expression atlas, both C_4_ decarboxylase genes decreased in relative expression as the leaf aged (Figure 3B). For both genes, expression was highest in the youngest leaf tissues and decreased continually in older leaves; the trend held for both the leaf at the top of the canopy and the leaf subtending the ear (Figure 3B). On average, NADP-ME3 was expressed at a level 2.4 times higher than PEPCK1. Unlike in the gene expression atlas, PEPCK1 expression was less divergent fromNADP-ME3 expression in the youngest leaves, and more divergent in older leaves. Leaf position significantly impacted the expression of both NADP-ME3 and PEPCK1 expression, but not the ratio between the two genes.

Decarboxylase activity was also measured using enzyme activity assays. Maize plants exhibited variation in both the amount of enzyme activity and the ratio of NADP-ME to PEPCK activity depending on both canopy position of the leaf and growth stage of individual leaves (ANOVA activity ~ leaf * plant age, p < 0.05). Both enzyme activities in the top leaf were not significantly different with plant age. Canopy position had a large effect, with leaves in the top of the canopy exhibiting 35.7% more PEPCK and 62.2% more NADP-ME activity in the four growth stages where both leaves were measured (Figure 4). In the top of the canopy, NADP-ME activity was 2-fold higher than PEPCK on average; however, in the subtending leaves, the ratio decreased to 1.5-fold higher, indicating a decreased relative role for NADP-ME in that part of the canopy. Thus, in the top leaves, 66% of decarboxylase activity came from NADP-ME and 34% from PEPCK, while in the subtending leaves, the proportion shifted to 60% from NADP-ME and 40% from PEPCK.

**Figure 4:**
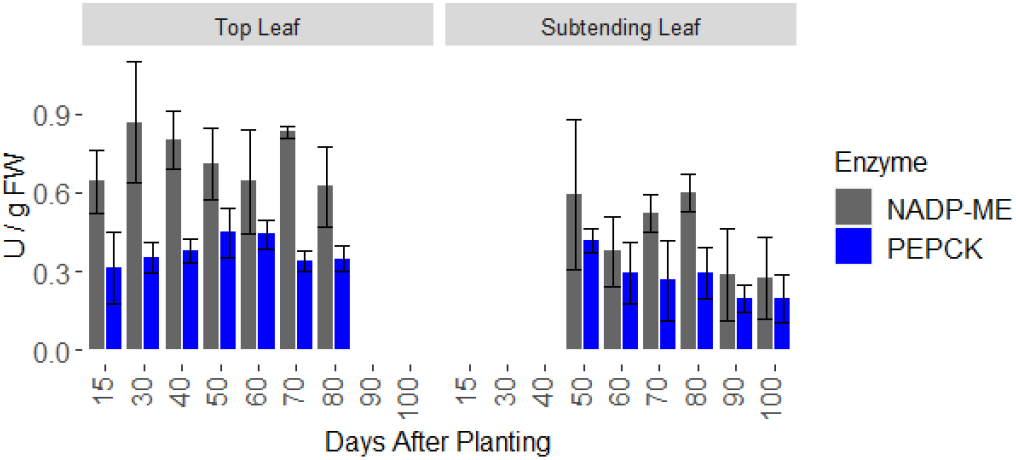
Enzyme activity assays: *In vitro* activities of NADP-ME (gray) and PEPCK (blue) from whole leaf segments of maize leaves at nine growth stages from the top or subtending leaf. Data are means ± sd of 3-4 replicate plants.

### Pool sizes of C_4_ and C_3_ metabolites

Malate and aspartate are the transfer acids that move CO_2_ from the mesophyll to the bundle sheath. Concentrations of both metabolites were measured in the two leaves for nine growth stages with LC-MS/MS. In the 15-day old plants, the ratio of malate to aspartate was 4.6:1, similar to reported values (Chapman and Hatch, 1981; Weissmann et al., 2016; Arrivault et al., 2017) and with expected pool sizes: 6.5±1.4 μmol malate g^−1^ FW and 1.4±0.6 μmol aspartate g^−1^ FW (Khamis et al., 1992; Lohaus et al., 1998; Szecowka et al., 2013; Arrivault et al., 2017). Both plant age and canopy position had significant effects on the pool sizes of malate and aspartate (ANOVA, type II, p<0.0001; Figure 5). In the top leaf, the pool size of malate decreased from 6.5±1.4 μmol g^−1^ FW at 15 days to 2.2±0.8 μmol g^−1^ FW for 30-80 DAS, while the aspartate pool size was largely unchanged with time. Thus, the ratio between the malate and aspartate pools decreased to 1.8±0.4 in the top leaf from 30 to 80 DAS (Figure 5). In the subtending leaf, the concentration of malate was generally greater than in the top leaf and both pool sizes decreased with plant age (Figure 5). In contrast, metabolites involved in the Calvin Benson Cycle were significantly affected by plant age, but only GAP/DHAP and E4P were significantly different between parts of the canopy, both having larger pools in the subtending leaf (Supplemental Figure 1).

**Figure 5:**
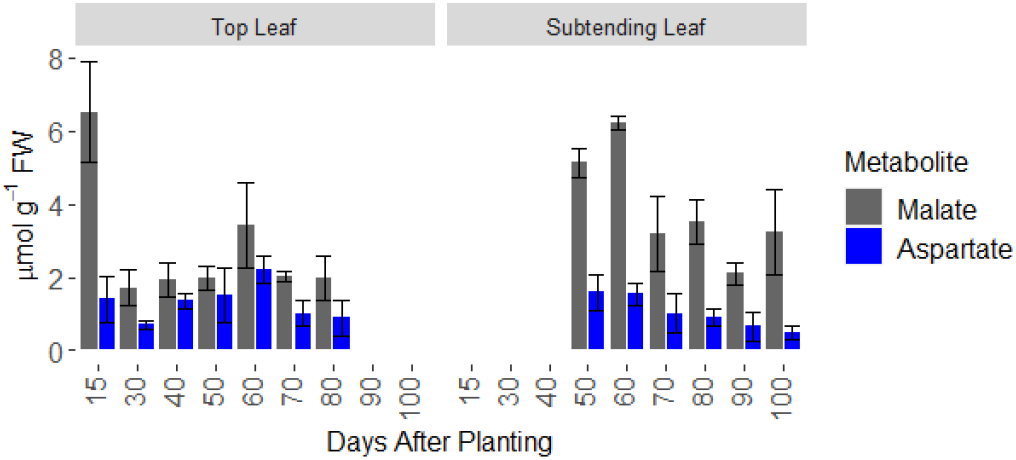
C_4_ transfer acid quantification: Malate and aspartate pools were quantified using an LC-MS approach and normalized by sample fresh weight. Malate pools (black) were larger than aspartate pools (blue) for all measured time points. Data are means ± sd of 3-4 replicate plants.

One hypothesis for why plants might use the NADP-ME and PEPCK pathway with similar ratios in subtending leaves and older leaves is to maintain smaller pools of the five transfer acids, rather than large pools of malate and pyruvate needed for the NADP-ME pathway alone. Pyruvate, PEP, and aspartate all had strong age effects, decreasing with leaf age, while alanine increased in the top leaf with plant age, and the malate pool size did not change with plant age. Because the malate pool is mostly inactive (discussed below), its pool was excluded from a total transfer acid pool comparison. This combined pool was slightly larger in the top leaf and smaller in the bottom leaf (p=0.055, ANOVA type II), and decreased with leaf age (p=1.56e-5, ANOVA type II)

### ^13^CO_2_ labeling in C_3_ and C_4_ metabolites

Differences in pool sizes for each growth stage may be the consequence of cumulative changes in metabolism over the life of the plant; however, isotopic labeling provides a snapshot of active metabolism. ^13^CO_2_ was provided to the leaf of 15-day or 55-day old plants, and metabolites were measured for label incorporation and absolute pool size. These two ages were chosen to represent plants at the beginning of the vegetative and reproductive growth stages, as well as contrasting C_4_ metabolite pool sizes. Plants were labeled for up to 5 minutes, and the average label incorporation in C_4_ shuttle metabolites and several central carbon metabolism metabolites were quantified (Figure 6A, Supplemental Figure 1). Leaves from 15-day old plants incorporated ^13^C label faster than the 55-day old plants, reaching an average labeling amount of 47-55% for measured Calvin Benson Cycle intermediates (i.e. phosphoglyceric acid (PGA), fructose bisphosphate (FBP), glyceraldehyde 3-phohsphate (GAP) and dihydroxyacetone phosphate (DHAP)) at 15 days compared to 12-27% for 55-day old plants in the top or subtending leaf, in agreement with the net CO_2_ assimilation data (Figure 1). The C_4_ intermediates were less enriched at both ages: 15-day old plants contained pyruvate that was 20% average labeled, 24.6% labeled phosphoenolpyruvate (PEP), 35% labeled aspartate, and 8.5% labeled malate. In the 55-day old plants, C_4_ intermediates were approximately half the average label of the 15-day old plants after 5 minutes of exposure to ^13^CO_2_ (Supplemental Figure 1).

**Figure 6:**
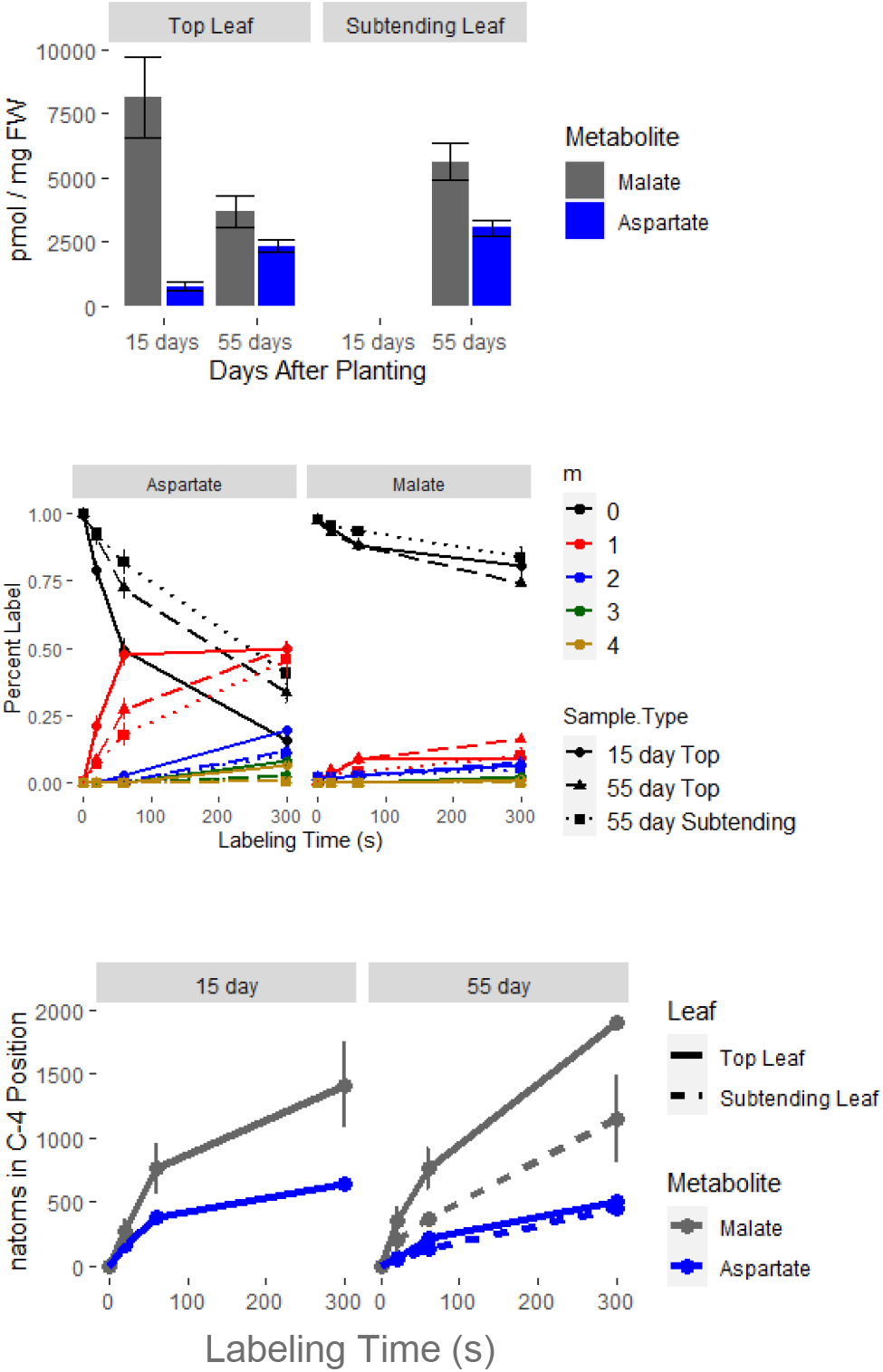
C_4_ transfer acid labeling: A.) Quantitation of malate and aspartate pools using LC-MS. Malate (gray); aspartate (blue). B.) Time course of labeling for malate and aspartate. Mass isotopolgs (m_n_) represent malate or aspartate molecules that have incorporated n molecules of ^13^C. C.) Moles of ^13^C in the C-4 position of malate (gray) and aspartate (blue) using the method of (Arrivault et al., 2017).

Malate and aspartate predominantly incorporate label from ^13^CO_2_ in the C-4 position, which is subsequently decarboxylated in the bundle sheath causing loss of the label (Chapman and Hatch, 1981). Label incorporation in the C1-C3 positions of malate and aspartate comes from downstream PEP and pyruvate labeling as a result of Calvin Benson Cycle activity. The malate pool also accumulates little ^13^C label due to a large proportion of the malate pool not participating in photosynthetic metabolism, resulting in a large proportion considered an inactive pool during stable isotope labeling (Szecowka et al., 2013; Ma et al., 2014; Allen, 2016; Arrivault et al., 2017). The inactive pool in the 15-day and 55-day plants were similar, with approximately 80% of malate unlabeled after 5 minutes (Figure 6B) and only 10% average labeling (Supplemental Figure 1), though the final asymptotic value where M_0_ levels off is incompletely defined at the five min time point (Arrivault et al., 2017; AuBuchon-Elder et al., 2020). Looking more deeply at the distribution of isotopologues, malate and aspartate labeling matched expectations based on the C_4_ pathways (Figure 6B). The m+1 isotopologue for aspartate increased rapidly from 0 to 60 seconds to account for 50% of all aspartate in 15-day old plants and 20-30% in subtending and top leaves of 55-day old plants. Malate m+1 isotopologues accounted for 5-10% of all malate. Very little (<5%) label accumulated in the C1-C3 positions of malate and aspartate during the first minute of labeling, also consistent with C_4_ topology.

The metabolism of the NADP-ME and PEPCK pathways was calculated in a cross-comparable way in the three leaves (15-day top, 55-day top, 55-day subtending) using the method described in Arrivault et al (Arrivault et al., 2017) to calculate n-atom equivalents in the C-4 position of malate and aspartate. N-atom equivalent measurements for malate and aspartate use the number of labeled carbons, the metabolite concentration, and the assumption that all malate or aspartate molecules with at least one ^13^C contain label in the C-4 position to quantify labeling from a molar basis (see Supplemental Table 1). From this calculation, malate and aspartate labeling were directly compared. During the first 60 seconds of labeling, the C-4 position of malate labeled at twice the rate of aspartate in the 15-day old plants and 4.3 and 2.6 times faster in the 55-day old top leaf and subtending leaf, respectively (Figure 6B), reflecting 66.5% of C_4_ activity going through the NADP-ME pathway and 33.5% through PEPCK and indicating a larger role for aspartate than previously identified, where aspartate only accounted for 10-25% of the activity of the C_4_ shuttle (Chapman and Hatch, 1981; Weissmann et al., 2016; Arrivault et al., 2017). In the 55-day old plants, the rate of labeling in the C-4 position of malate in the subtending leaf was approximately one-half the top leaf, while aspartate labeling was consistent throughout the canopy, resulting in 18.8% of C_4_ activity through PEPCK in the top leaf, and 27.9% in the subtending leaf (Figure 6C).

### Asparagine concentrations in the plant

Asparagine is an important storage and transport form of nitrogen in maize plants (Lohaus et al., 1998; Lea and Azevedo, 2007) and has been shown to specifically induce PEPCK activity in grape seeds (Walker et al., 1999). Asparagine can be synthesized from aspartate via asparagine synthetase and converted back to aspartate through L-asparaginase; however, the relationship between the C_4_ PEPCK and asparagine concentration is not known.

Asparagine concentration was measured using LC-MS/MS. The asparagine concentration in the leaf varied from 0.15 μmol mg^−1^ to 6.6 μmol mg^−1^ and did not have a clear pattern with leaf age or canopy position in this experiment (Figure 7). Concentrations were more variable than other metabolites (Supplemental Figure 1), resulting in a decreased ability to observe any trends in the data. As such, asparagine was very weakly correlated with PEPCK activity (Pearson’s R = 0.17) and somewhat correlated with PEPCK expression (Pearson’s R = 0.32) in the data.

**Figure 7:**
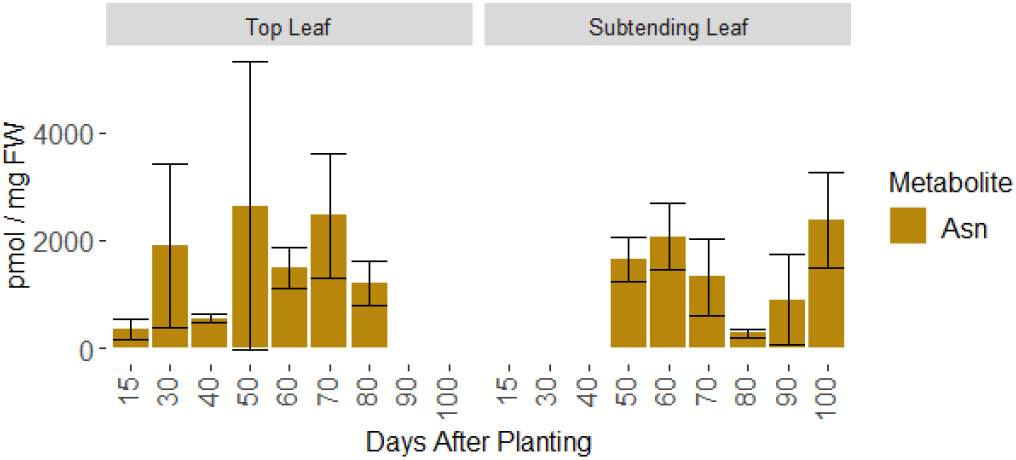
Asparagine: asparagine concentration during plant growth measured by LC-MS normalized by sample fresh weight. Data are means ± sd of 3-4 replicate plants.

PEPCK activity was more closely correlated with its substrate, aspartate (R=0.74), although it was not correlated with PEPCK expression (R=0.07). Aspartate and asparagine correlations were not correlated (R = 0.01). No correlation was observed between asparagine concentration and PEPCK expression or activity, predominantly because asparagine concentrations were highly variable within samples, but similar in all plant ages and both canopy positions. To fully elucidate this relationship in maize leaves, a more targeted approach would be necessary, focusing on asparagine concentrations in bundle sheath cells. Moreover, asparagine concentration fluctuates diurnally, with its highest concentrations at night (Harmer et al., 2018; Kambhampati et al., 2018), so the relationship between asparagine concentration and PEPCK activity in the leaf may also have a diurnal component which was not assessed in these experiments.

## Discussion

The activities of C_4_ subtype pathways vary during maize growth; with the greatest difference between leaves at the top and middle of the canopy, rather than with plant age. PEPCK activity was consistent between the top leaf and subtending leaf; however, NADP-ME activity decreased in the older subtending leaf compared to the top leaf, leading to a decreased overall rate of decarboxylation and changing the ratio of the decarboxylases between the two canopy positions. Similarly, the PEPCK pathway metabolite aspartate labeled at the same rate in the top and subtending leaf of 55-day old plants and correlated with PEPCK activity, while malate labeling decreased by three-fold in the subtending leaf compared to the top leaf. Decreased malate labeling in the subtending leaf was surprising considering the subtending leaves from all plant growth stages had larger malate pools than top leaves. These results indicate a large, inactive pool of malate in the subtending leaf, in addition to the photosynthetically active pool of malate in plant leaves (Szecowka et al., 2013; Ma et al., 2014; Weissmann et al., 2016; Arrivault et al., 2017). The large malate pool may serve to buffer the fluctuating light environment within the canopy or could function as a carbon reserve for remobilization to the grain.

Fluctuating environmental conditions, in particular with regard to light, are hypothesized to be a reason that plants might use a combination of C_4_ pathways in parallel (Furbank, 2011; Bellasio and Griffiths, 2014a; Stitt and Zhu, 2014; Wang et al., 2014). The more stable, typically unshaded light environment at the top of the canopy may not require the same degree of metabolic flexibility afforded by parallel pathways and benefits from the efficiency of the NADP-ME pathway with CO_2_ decarboxylation directly in the chloroplast (Wang et al., 2014). The subtending leaves receive less light in a closed canopy but can have high intensity sun flecks under field conditions and thus may benefit through shared C_4_ pathway operation (Bellasio and Griffiths, 2014a; Stitt and Zhu, 2014; Wang et al., 2014). Our experiments suggest the balanced activities occur though a reduction in NADP-ME activity and decreased total C_4_ shuttling in lower parts of the canopy (Figure 4, Figure 6), which did not result in a lower carbon assimilation rate or reduced Calvin Benson Cycle metabolite labeling. Carbon assimilation rates were however significantly affected by the age of the plant (Figure 2, Supplemental Figure 1). Subtending leaves maintained the same level of CBC metabolism as leaves at the top of the canopy, likely as a result of even light distribution through the canopy due to growth in a greenhouse rather than a dense field canopy (Collison et al., 2020).

This study was designed to address differences in C_4_ metabolism based on leaf age and canopy position, such that the same two leaves were compared between plants of different ages. There were few differences in C_4_ metabolism between plants of different ages (i.e. top leaf on plants), but significant differences in C_4_ pathways between leaves in different positions on the same plant (i.e. microenvironment). These results bring up important possibilities of regulation by light that differ between leaves on the same plant. Light may play a crucial role in regulating the C_4_ decarboxylases between canopy positions. The relationship between light and C_4_ pathway activity has been studied in the context of different species with different C_4_ subtypes (Ubierna et al., 2013; Sonawane et al., 2018). Less consideration has been given to the plasticity of the C_4_ pathways within a species. NADP-ME is strongly light regulated by light quality (Casati et al., 1998) and quantity (Hatch and Kagawa, 1976; Bellasio and Griffiths, 2014b), and by time of day, through phosphorylation, with its activity peaking at two hours after dawn and decreasing slowly throughout the day (Bovdilova et al., 2019). Leaves positioned lower in the canopy may not receive the quality and quantity of light to activate NADP-ME to the levels observed at the top of the canopy (Figure 4, Figure 6). In contrast, PEPCK activity may be unresponsive to light (Wingler et al., 1999) or potentially negatively regulated by light (Chao et al., 2014); though it should be noted that the experiments reported here were performed in the greenhouse where plant leaves are not shaded to the extent they would be in the field. PEPCK expression levels are more clearly responsive to other environmental conditions such as N supply which could have effects in leaves subtending the ear (Delgado-Alvarado et al., 2007; Penfield et al., 2012). N distribution in the canopy also regulates the level of photosynthesis in herbaceous canopies (Niinemets, 2016).

## Conclusions

The top and middle of the canopy and different ages of the plant experience changes in C_3_ and C_4_ cycle metabolism. Variation in the relative proportion of the two C_4_ pathways was strongest between leaves at the top and bottom of the canopy, while differences in overall photosynthetic rate were predominantly caused by the age of the plant. C_4_ characteristics changed as the leaf aged, although they did not shift relative to each other, instead generally decreasing overall to coordinate with decreased C_3_ metabolism. At the top of the canopy, the plant predominantly uses NADP-ME and shifts to more equal use of the two C_4_ pathways lower in the canopy, as the microenvironment in the lower canopy is more variable and the leaves shift to supporting growth of sink tissues throughout the plant. Plasticity within the plant life cycle and within the canopy is a potential advantage of the parallel C_4_ pathways, allowing fine-tuning of the C_4_ pathways to optimize for specific growth conditions of the leaf.

## Methods

### Plant Growth

Plants were grown in the greenhouse in Saint Louis, MO from February to July 2019. Four maize plants (genotype W22) were planted at ten day intervals for 90 days. Plants were self-pollinated upon flowering. Plants were grown in Berger 35, 7% bark medium in 2.5 gallon pots. Plants were grown with 14-hour day length, 10 hour nights using supplemental light to extend the day length and when available sunlight was below 600 μmol m⁻² s⁻¹. Plant growth temperature was 28°C day, 22°C night and a minimum 40% relative humidity.

Plants were sampled at 100 days after initiation of the experiment. Plants were 10, 20, 30, 40, 50, 60, 70, 80, or 90 days old at the time of sampling. Leaf tissue was collected from the topmost leaf and the leaf subtending the ear (leaf 13 in W22), when available. For the two oldest sets of plants (80 or 90 days after sowing), the topmost leaf on each plant was fully senesced and not sampled. For the three youngest sets of plants (10, 20, or 30 days after sowing), the subtending leaf had not emerged from the whorl and was not sampled. All measurements were taken from 10cm leaf segments beginning 20cm from the tip of the leaf.

### Gas Exchange Measurements

Net CO_2_ assimilation was measured using gas exchange measurements on a LI-6800 Photosynthesis System (Li-Cor Inc., Lincoln, Nebraska). Net CO_2_ assimilation was measured at steady state at 400ppm CO_2_ and 350 followed by 1500 μmol m⁻² s⁻¹ light. The leaf was clamped into the instrument head using a 6cm aperture, temperature was controlled at 25°C, and the gas flow rate was 500 μmol s^−1^.

### Gene Expression of PEPCK and NADP-ME

Leaf tissue was sampled from a 10cm segment starting 20cm from the leaf tip, from either the leaf at the top of the canopy or from leaf number 13, which is the leaf below the ear in W22 (subtending leaf). RNA was extracted from 50mg of leaf tissue using Trizol following the manufacturer instructions (Invitrogen 15596026). Residual genomic DNA was removed using DNAseI (Invitrogen TURBO DNA-free, AM1907), and first strand cDNA synthesis was performed using Invitrogen Superscript II (18064022). Gene expression for PEPCK1 (Zm00004b001002) and NADP-ME3 (Zm00004b015828) were quantified via qRT-PCR using a Roche Light Cycler 480 II using Roche SYBR green I (Roche 04707516001). The delta cT method was used to quantify relative gene expression of PEPCK1 and NADP-ME3 over time. NAC26 (Zm00004b022707) was used as a housekeeping gene using published primers (CCGCCGTCAACAGGGAAATCTG, GTAGCACGCCCAAGACCAACAG; (Lin et al., 2014)Genome-wide identification of housekeeping genes in maize). Primers for PEPCK1 were forward: CCCGATCAACACCTGGACG and reverse: GACGCACCCATGACAATACC; primers for NADP-ME3 were forward: GAGTCAGGGCCGTTCAATCT and reverse: ACAGAGTACCATCCGCGTTG.

### Enzyme Activity

PEPCK activity was measured using the method of Walker et al (Walker et al., 1999) detailed on protocols.org (Osorio et al., 2014). Briefly, crude protein was extracted from ~100mg of fresh leaf tissue in a buffer containing 0.5M bicine-KOH (pH 9.0), 0.2M KCl, 3mM EDTA, 5% (w/v) PEG-4000, 25mM DTT and 0.4% bovine serum albumin. The extract was centrifuged for 20 minutes and the supernatant was added to a buffer containing 0.5M bicine-KOH (pH 9.0), 3mM EDTA, 55% w/v) PEG-4000, and 25mM DTT. The sample was incubated for 10 minutes on ice, centrifuged at 13,000 x g at 4°C for 20 minutes. The supernatant was discarded and the pellet was resuspended in 10mM bicine-KOH (pH 9.0) with 25mM DTT. Activity was measured in the carboxylation direction by coupling the reaction with malate dehydrogenase and following the oxidation of NADH at 340nm using Molecular Devices SpectraMax M2 spectrophotometer. Total protein was measured using the protein extract for PEPCK activity using Bradford reagent (Millipore Sigma; Cat: B6916) and commercial bovine serum albumin standards (Thermo Fisher 23208). NADP-malic enzyme activity was measured using the method described in Osorio et al (Osorio et al., 2014) from Detarsio et al (Detarsio et al., 2003), using 50mg of fresh leaf tissue. Enzyme activity was measured by following the reduction of NADP^+^ at 340nm for 5 minutes.

### Compositional Analysis

Chlorophyll content was measured using the method of Arnon et al (Arnon, 1949). Amino acid, sugar and sugar phosphate content was measured using the method described in Czjaka et al, (Czajka et al., 2020). Briefly, metabolites were extracted from 100mg fresh weight of ground leaf tissue using 3:7 (v/v) methanol:chloroform solution incubated for two hours on a rotator at 4°C. After two hours, 0.5 mL of ddH_2_O was added, the solution was centrifuged and the supernatant (aqueous phase) was centrifuged in 3KDa filters, frozen, lyophilized, and resuspended in 50uL 1:1 methanol:ddH_2_O. Metabolite quantities were measured using LC-MS using a Shimadzu Prominence-xR UFLC system and a SCIEX hybrid triple quadrupole-linear ion trap MS equipped with Turbo V™ electrospray ionization source for separation and detection of metabolites, and samples were injected into an InfinityLab Poroshell 120 HILIC-Z (2.1 x 100 mm, 2.7 μm, Agilent Technologies) column.

### Stable isotope labeling using ^13^CO_2_

Isotopic labeling was performed on plants from two of the stages, 15 days and 55 days after sowing. For 15-day old plants, the top collared leaf was sampled, and for 55-day old plants, both the top collared leaf and the subtending leaf were used. The 15-day old plants were selected to represent the growth stage commonly characterized for C_4_ pathway labeling (Chapman and Hatch, 1981; Weissmann et al., 2016; Arrivault et al., 2017), where young maize leaves use the NADP-ME pathway for 75-90% of metabolism, relying on the PEPCK pathway as a minor contributor. The 55-day sample was selected to represent plants that have generated all of their leaves, reached the end of vegetative growth, and are approaching the developmental shift at anthesis. Labeling in central carbon metabolites was measured using ^13^CO_2_ provided to the leaf with a hand-held clamp for 0, 20, 60, or 300 seconds to generate a time course of label incorporation. Leaves were freeze clamped at the end of each time interval, flash frozen in liquid nitrogen, and stored at −80°C until further processing. Metabolites were extracted in chloroform:methanol as above, and metabolite labeling and pool size were measured by LC-MS/MS on a Qtrap6500 (Chu et al, in prep; (Czajka et al., 2020)). Samples were corrected for natural abundance of ^13^C using IsoCorrectoR (Heinrich et al., 2018).

### Statistical Analysis

ANOVA comparisons were performed using two-way ANOVAs with factors canopy position (leaf) and plant age, with the Anova function in the car package in R (Fox and Weisberg, 2018) to use a type II ANOVA to be able to compare factors with unequal sample sizes.

## Acknowledgements

The authors would like to thank the Donald Danforth Plant Science Center Plant Growth Facility and Donald Danforth Plant Science Center Proteomics and Mass Spectrometry Facility. Thank you to Emma Smith, Emily Frankenreiter, and Genesis Hudson for assistance with performing tissue sampling.

## Supplemental Data

Supplemental Table 1: Calculations to determine degree of labeling in the C-4 position of malate and aspartate.

Supplemental Figure 1: A.) Metabolite quantitation of C_4_ intermediates. PEP: phospho*enol*pyruvate. B.) Metabolite quantitation of Calvin Benson Cycle intermediates which were able to be quantified. TP: Triose phosphate (i.e. GAP and DHAP: Glyceraldehyde 3-phosphate and Dihydroxyacetone phosphate), FBP: Fructose 1,6-bisphosphate; E4P: Erythrose 4-phosphate; S7P: Sedoheptulose 7-phosphate; R5P: Ribose 5-phosphate.

Supplemental Figure 2: Average labeling in C_3_ and C_4_ metabolites during the five minute time course. PEP: phospho*enol*pyruvate, PGA: phosphoglyceric acid, FBP: fructose 1,6-bisphosphate, TP: triose phosphate (i.e. GAP and DHAP: glyceraldehyde 3-phosphate and dihydroxyacetone phosphate), UDPG: uridine diphosphate glucose, 2OG: 2-oxoglutarate, ASN: asparagine, GLN: glutamine, Glu: glutamate.

Supplemental Figure 3: Isotopologue distribution graphs for labeled metabolites during the five minute time course. PEP: phospho*enol*pyruvate, PGA: phosphoglyceric acid, FBP: fructose 1,6-bisphosphate, TP: triose phosphate (i.e. GAP and DHAP: glyceraldehyde 3-phosphate and dihydroxyacetone phosphate), UDPG: uridine diphosphate glucose, 2OG: 2-oxoglutarate, ASN: asparagine, GLN: glutamine, Glu: glutamate.

